# Focused ultrasound programmed characteristic NIR-IIb lanthanide mechanoluminescence for high sensitivity bioimaging in vivo

**DOI:** 10.64898/2025.12.30.696464

**Authors:** Minglong Zhang, Jieying Zhang, Lingkai Meng, Xiaohe Li, Sixin Xu, Xingjun Zhu

## Abstract

Optical imaging techniques for biodetection are hindered by limited sensitivity and resolution in deep tissues, arising primarily from photon scattering and endogenous autofluorescence. Herein, we report a focused ultrasound (FUS) programmable NIR-IIb emissive mechanoluminescent nanoparticle for in vivo bioimaging. Specifically, CaZnOS nanoparticles co-doped with Mn^2+^ and Er^3+^ were engineered to exhibit mechanoluminescence (ML) emission peaked at 1550 nm. FUS stimulation elicits ML without the need for optical excitation, thereby enabling deep-tissue imaging devoid of background interference from excitation light. By modulating the excitation frequency of FUS, ML can be programmed as a characteristic signal. Integrated with a Fourier frequency-transform identification reconstruction (FFIR) algorithm, the specific ML signals can be detected with high sensitivity, reaching sub-millimeter spatial resolution and markedly enhanced signal-to-background ratios compared to conventional photoluminescence-based approaches. By macrophage internalization of the ML nanoparticles to yield functional macrophage-nanoparticle (FMNs), a ultralow detection limit of 10 cells can be achieved. In murine tumor models, the FMNs facilitated real-time monitoring of early-stage tumor progression, underscoring their biocompatibility and translational promise for high-precision diagnostics in deep-seated neoplasms.

## 1. Introduction

Optical imaging (OI) has emerged as a promising technology for tumor detection, leveraging its capacity for real-time visualization, relative ease of molecular targeting, and lack of ionizing radiation^1-3^. However, the penetration depth and spatial resolution of OI in the visible (400-700 nm) and the first near-infrared (NIR-I, 700-900 nm) windows are substantially limited by strong tissue absorption and scattering^4, 5^. Imaging in the second near-infrared window (NIR-II, 1000-1700 nm) mitigates this issue by reducing scattering, thereby improving deep-tissue imaging quality^4, 6, 7^. Nevertheless, current NIR-II luminescence imaging relies predominantly on photoluminescent probes, whose sensitivity remains compromised by exogenous laser excitation and background autofluorescence^8^. These limitations restrict applications in trace-level tracking and fine-structure imaging. Consequently, the precise observation of tumor microstructure and dynamic monitoring of pathological states in deep-seated solid tumors still lack high-resolution, high-sensitivity, and microenvironment-specific imaging technologies^9^.

Recent advances in mechanoluminescence (ML) materials offer a novel strategy to address these challenges. Upon mechanical stimulation, such materials develop an internal piezoelectric field via force-induced charge separation. Subsequent carrier-hole recombination after force removal activates luminescent centers, producing photon emission. Crucially, ML operates without optical excitation^10, 11^, effectively eliminating the background interference inherent in conventional photoluminescence. Moreover, by harnessing multi-level energy transfer among ions, ML enables multimodal light conversion and tunable emission properties^11-16^. Conventional ML materials (e.g., ZnS), however, often exhibit short emission wavelengths and weak intensity, hindering biomedical use^11^. In contrast, CaZnOS (CZOS), a wide-bandgap semiconductor ML host, shows considerable promise owing to its excellent stability and tunable luminescence^13, 17-20^.

Lanthanide doping in CZOS represents a feasible route for generating NIR luminescence. Simultaneous co-doping of lanthanide ions with transition metal ions, such as manganese ions (Mn^2+^), can enhance luminescence efficiency. Lei et al. confirmed that Mn^2+^, acting as a sensitizer, significantly enhances the emission of Nd^3+^-doped CZOS in the 900-1200 nm range^21^. It has been reported that under mechanical force, the bending of the valence and conduction bands in CZOS:Mn,Ln materials due to lattice strain triggers charge separation and rapid recovery processes, releasing energy that is rapidly transferred to Mn^2+^. Serving as an energy transfer hub, energy is efficiently transferred from Mn^2+^ to Ln^3+^ via a resonant energy transfer mechanism. Concurrently, Mn^2+^ doping narrows the host bandgap, reduces the energy level mismatch between host electron-hole recombination and Ln^3+^ excited states, promoting further energy transfer to Ln^3+^and therey significantly enhancing emission^19, 21^.

Based on these advancements, the lanthanide mechanoluminescence intensity of CZOS in NIR-IIb window was optimized through a Mn^2+^ and Er^3+^ co-doping strategy. The hydrophilic modified nanoprobes are used to label tumor-associated macrophages (TAMs), thereby constructing a living cell-based biomimetic luminescent sensor (FMN). Leveraging the homing capability of macrophages to the tumor neovascular microenvironment, the probes target and label early-stage tumor regions with high proliferative activity, providing precise diagnostic results on pathological status to guide treatment. Focused ultrasound (FUS), as a non-invasive, highly spatially selective external modulation technique, can penetrate micro-scale biological tissues and generate mechanical stress in targeted areas, offering unique advantages for deep-tissue imaging^22, 23^. FUS was utilized to directly excite NIR-II emission of FMNs without external lasers. Simultaneously, by modulating the excitation frequency of FUS and combining it with the Fourier Frequency-transform Identification Reconstruction (FFIR) algorithm developed for frequency screening, it was possible to achieve highly sensitive and high-resolution tracing on a minimum of 10 FMNs with sub-millimeter spatial resolution, offering a novel for precise diagnosis of deep-seated tumors (Fig. 1).

**Figure 1.**
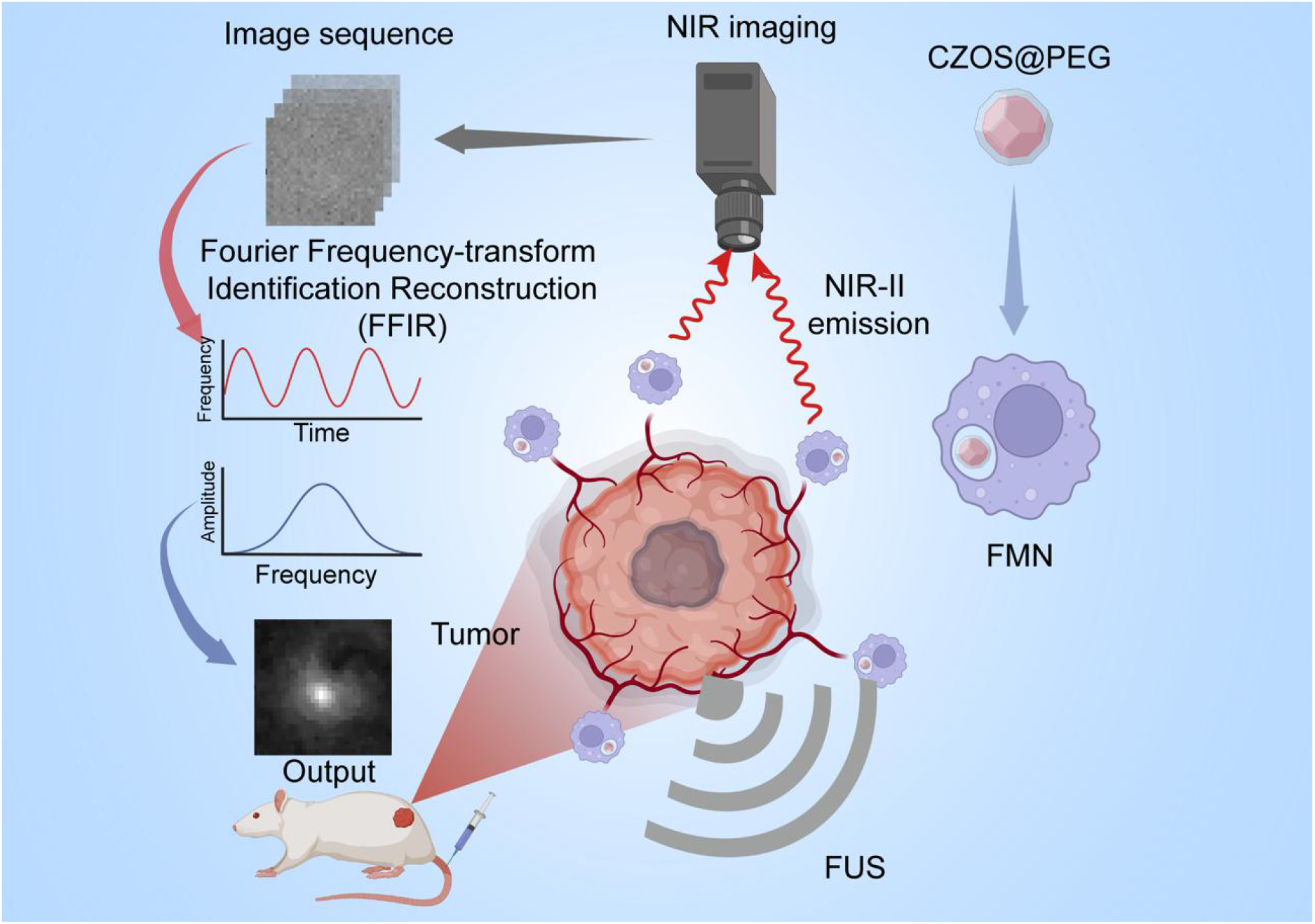
Schematic illustration of the focused ultrasound (FUS)-responsive high-sensitivity near-infrared-II (NIR-II) imaging system for monitoring the migration of macrophages at early stage of tumors.

## 2. Results and discussion

### 2.1 Synthesis Optimization and Characterization of CZOS

To optimize the luminescence intensity of Ca^1-y^Zn^1-x^OS: xMn, yEr (CZOS) in the NIR-II window, a series of samples with different Mn^2+^ and Er^3+^ doping concentrations were prepared and their photoluminescence (PL) properties were compared to determine the optimal doping ratio. Referring to the precursor concentrations previously used by Zhang et al.^19^, the Er^3+^ doping concentration was first varied while maintaining a fixed 2% Mn^2+^ doping level to observe changes in the luminescence intensity of CZOS in the visible and near-infrared windows. Upon irradiation with a continuous-wave (CW) 365 nm laser, the luminescence intensities of CZOS at both 610 nm and 1550 nm initially increased and then decreased with increasing Er^3+^ concentration, reaching a peak at an Er^3+^ doping concentration of 6% (Fig. 2a and 2b). Based on this, the Mn^2+^ doping concentration was subsequently adjusted while keeping the Er^3+^ concentration constant at 6%. Under the same UV-PL conditions, the luminescence intensity at 610 nm gradually decreased with increasing Mn^2+^ concentration, accompanied by a slight blue shift in the emission peak position. Concurrently, the emission at 1550 nm initially increased and then decreased, reaching its maximum intensity at a Mn^2+^ doping concentration of 50% (Fig. 2d and 2e). This phenomenon suggests that in CZOS, low Mn^2+^ concentrations cannot effectively sensitize the 1550 nm emission of Er^3+^. As the Mn^2+^ concentration increases, the energy migration (EM) process between Mn^2+^ and Er^3+^ gradually strengthens, promoting the Er^3+^ transition from the ^4^I_13/2_ to ^4^I_15/2_ energy level, manifested as enhanced luminescence at 1550 nm.

**Figure 3.2.**
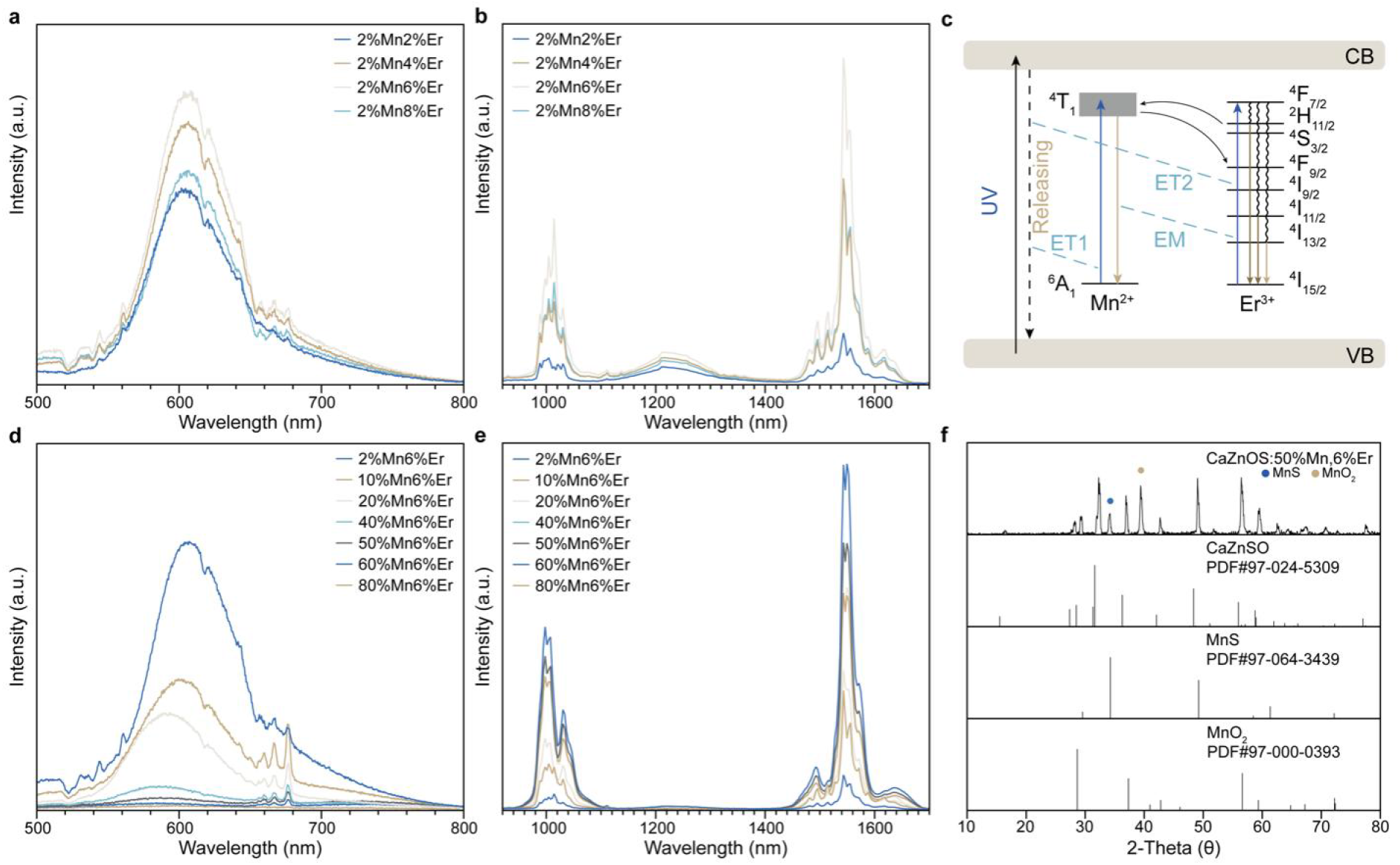
Luminescent spectra of CaZnOS:2%Mn, yEr in (a) visible window and (b) near infrared window. (c) Schematic diagram of the photoluminescence (PL) mechanism in CZOS. Luminescent spectra of CaZnOS:xMn, 6%Er in (d) visible window and (e) near infrared luminescent spectra of CaZnOS:xMn, 6%Er. CaZnOS:xMn, yEr was excited by a continuous-wave (CW) 365 nm laser and a 400 nm long-pass filter was used to exclude excitation. (f) X-ray powder diffraction (XRD) patterns of CaZnOS:50%Mn, 6%Er (CZOS).

However, with a continued increase in Mn^2+^ concentration, the EM between Mn^2+^ and material defects gradually intensifies, hindering further energy transfer to Er^3+^, resulting in a decrease in the 1550 nm emission intensity after reaching its peak (Fig. 2c). Based on the requirement for high luminescence efficiency in the NIR-II window, the optimal doping ratio for CZOS was ultimately determined to be 6% Er^3+^ and 50% Mn^2+^. X-ray powder diffraction (XRD) results revealed that CZOS doped with 6% Er^3+^ and 50% Mn^2+^ exhibited characteristic peaks corresponding to three crystalline phases: CaZnSO (PDF#97-024-5309), MnS (PDF#97-064-3439), and MnO_2_ (PDF#97-000-0393) (Fig. 2f).

### 2.2 Verification of Ultrasound-Induced Mechanoluminescence (USML) of CZOS

Subsequently, the USML performance of CZOS was characterized using an encapsulated phantom. The luminescence mechanism is illustrated in Fig. 3a ^19^. When the CZOS phantom was exposed to FUS (repetition frequency: 1 Hz, duty cycle: 15%, power: 24.15 W cm^-2^), a flickering spectral signal was captured by a spectrometer within the NIR-II window. This signal exhibited a peak wavelength consistent with the previously observed UV-PL signal (Fig. 3.5b), confirming that CZOS can effectively respond to FUS excitation to produce emission at 1550 nm. Under constant ultrasound excitation parameters (repetition frequency: 1 Hz, duty cycle: 15%, power: 10.88 W cm^-2^), the CZOS phantom underwent multiple cycles of ultrasound treatment. Observation was conducted using an imaging system equipped with an InGaAs camera, collecting images continuously with an exposure time of 100 ms. Analysis of the resulting image sequence revealed that CZOS could be stably excited by ultrasound to produce pulsed luminescence without requiring additional optical pre-charging. The luminescence intensity remained consistent across 10 excitation cycles (Fig. 3.5c), demonstrating excellent repeatability of the ML generated by FUS excitation of CZOS.

**Figure 3.**
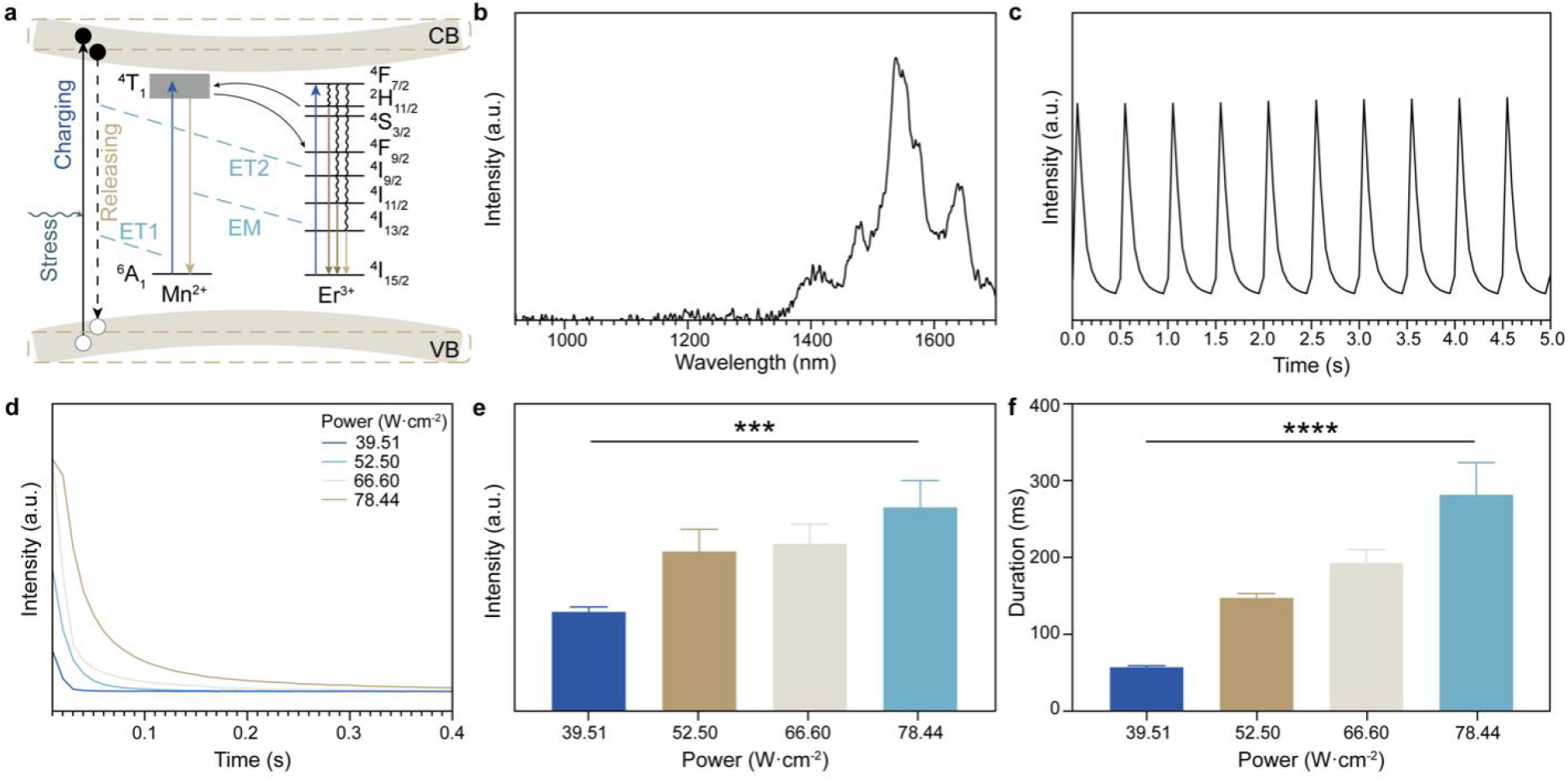
Near-infrared luminescence properties of CZOS under FUS excitation. (a) Schematic diagram of the ultrasound mechanoluminescence (USML) mechanism in CZOS. (b) NIR emission spectra of CZOS under FUS excitation. A1400 nm long-pass filter was used to exclude excitation. (c) NIR USML intensity of CZOS under multiple excitations. NIR USML (d) decay curve, (e) intensity and (f) duration of CZOS at different FUS power levels. A 1500 nm long-pass filter was used to exclude excitation when identifying the profiles of USML in CZOS.

Based on the reported ML mechanism^19^, CZOS, as a strongly piezoelectric wide-bandgap semiconductor, experiences a tilt in its conduction and valence band energy levels due to the internal piezoelectric potential when subjected to stress. This facilitates the detrapping of trapped electrons from defect states within CZOS into the conduction band. The internal ML intensity is expected to increase with the potential difference generated by the mechanical force. The detrapped electrons subsequently transfer energy (ET) to Mn^2+^ in its ground state, promoting it to the ^4^T_1_ energy level. Energy is then migrated (EM) to Er^3+^, inducing the ^4^I_13/2_ to ^4^I_15/2_ transition. Based on this, a positive correlation is hypothesized between FUS excitation power and both the USML intensity and lifetime of CZOS. To investigate this, a series of FUS power levels (39.51-78.44 W cm^-2^) were applied to sequentially excite the same CZOS phantom, and continuous images were collected with a 10 ms exposure time. Quantitative analysis of the luminescent regions in the images revealed that the peak USML intensity of CZOS in the NIR-II window increased significantly with higher FUS power (Fig. 3d and 3e). Concurrently, the USML lifetime also notably prolonged with increasing FUS power (Fig. 3d and 3f).

### 2.3 In Vitro Evaluation of High-Sensitivity Imaging Performance

Building upon the previously described FUS-responsive near-infrared luminescence imaging system, its feasibility for deep-tissue imaging was validated in vitro. CZOS was encapsulated within a cylindrical channel (diameter: 0.5 mm) using a hollow phantom model. Pulsed ultrasound (duty cycle: 15%) or laser illumination (808 nm, 0.5 W cm^-2^) was applied to the material-encapsulated region to evaluate the system’s imaging performance. To simulate biological tissue imaging conditions, a series of gelatin phantoms with varying thicknesses (0-5 mm) were placed atop the CZOS phantom to verify the system’s applicability at different tissue depths. Continuous images were collected with a 100 ms exposure time for quantitative analysis. Imaging results showed that the signal spot intensity for both PL and USML decreased with increasing tissue thickness. However, USML demonstrated superior descattering effects compared to PL, retaining more intensity information under equivalent tissue coverage, primarily due to its excitation-light-free imaging mechanism (Fig. 4a). After FFIR processing, the signal-to-background ratio (SBR) of the resulting USML images was significantly improved. Accurate visualization of the channel boundary remained possible until the overlying tissue thickness reached 5 mm (Fig. 4a).

**Figure 4.**
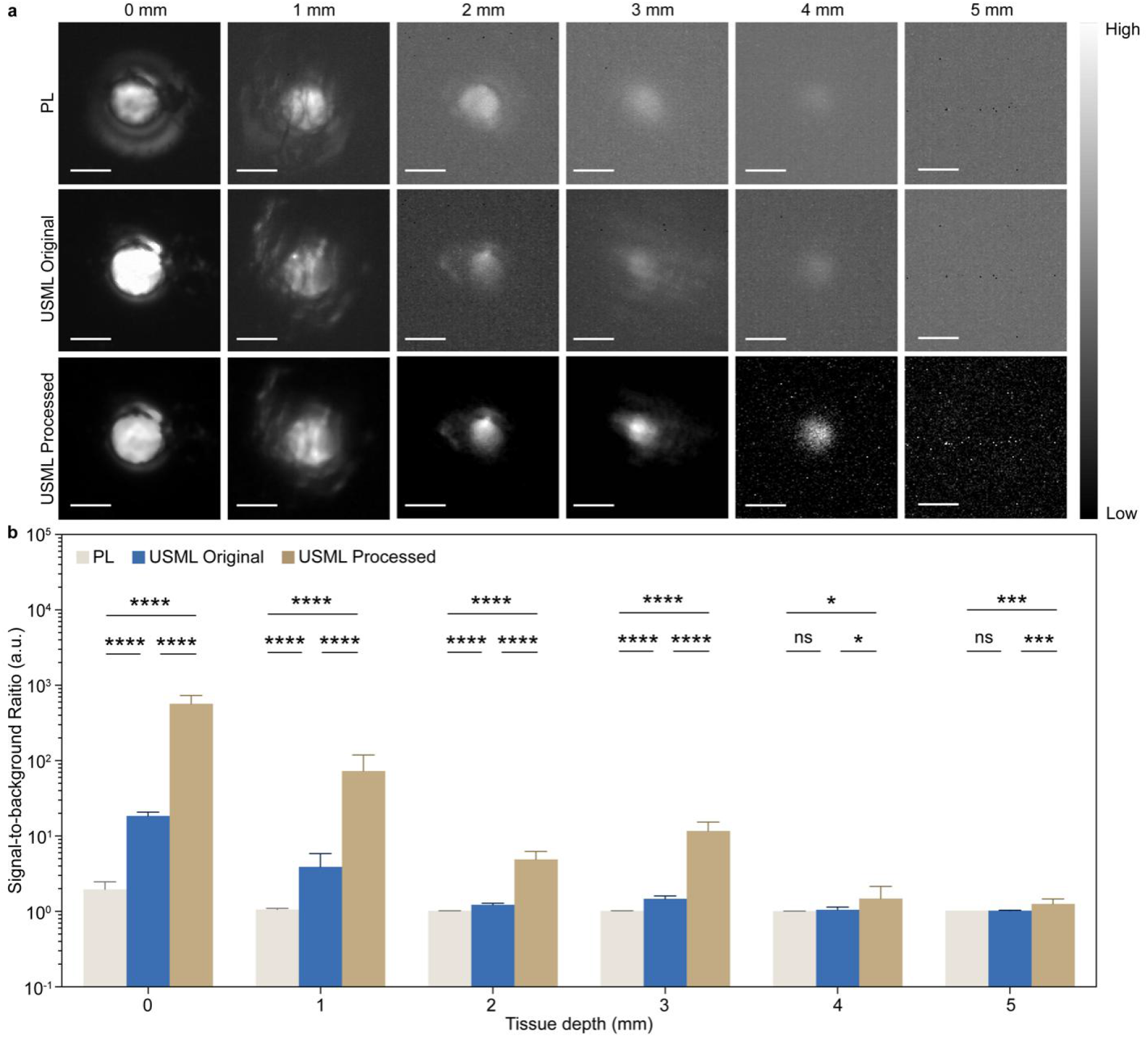
(a) PL images and ultrasound mechanoluminescence images images pre- and post-processing with the fourier transform-based frequency identification reconstruction (FFIR) algorithm at different tissue depths. All images have been normalized by intensity. (b) Signal-to-background ratio (SBR) comparison of PL imaging with USML imaging pre- and post-FFIR processing. Error bar stands for s. d. from n = 16 replicated samples. The scale bar was defined as 0.5 mm. FUS parameters applied to CZOS: repetition frequency = 1 Hz, duty ratio = 15%, and a 1500 nm long-pass filter was used to exclude excitation.

Further quantitative analysis revealed that without tissue coverage, the SBR for PL images was 1.93, while for USML images it was 18.28, representing a significant enhancement over PL. Moreover, after FFIR processing, the SBR of USML images could be further increased to 563.57, demonstrating a marked advantage in imaging resolution (Fig. 4b). When the phantom thickness reached 4 mm, PL imaging could barely locate the luminescent signal. In contrast, the luminescent characteristics of USML provided the prerequisite for FFIR processing. After FFIR optimization, the SBR could be increased to 1.46, presenting a relatively clear signal spot in the image. Based on these results, the excitation-light-free mechanism of USML grants it a significant advantage in imaging sensitivity for trace samples in deep tissue compared to PL. Concurrently, its frequency-specific luminescence characteristic also creates the prerequisite for FFIR processing, enabling further improvement in imaging quality.

To construct the living cell-based biomimetic probe, the existing CZOS required surface hydrophilic modification to enhance its biocompatibility and improve the efficiency of converting macrophages into the living cell probe (FMN). Grafting mPEG-Silane onto the CZOS surface yielded CZOS@PEG, which effectively slowed the hydrolysis rate of CZOS in aqueous environments and improved its intracellular stability. Transmission electron microscopy (TEM) imaging showed that the modified CZOS@PEG had a diameter of approximately 68.43 nm (Fig. 5a). Dynamic light scattering (DLS) analysis indicated that the particle size of CZOS increased from 72.07 nm to 86.35 nm after hydrophilic modification (Fig. 5b), while the Zeta potential increased from -0.17 mV to -24.83 mV (Fig. 5c), indicating improved hydrophilicity post-modification. Fourier-transform infrared spectroscopy (FTIR) analysis revealed characteristic peaks for C-N, N-H, and Si-O stretching vibrations, confirming the successful grafting of long-chain alkyl ligands onto the CZOS@PEG surface (Fig. 5d).

**Figure 5.**
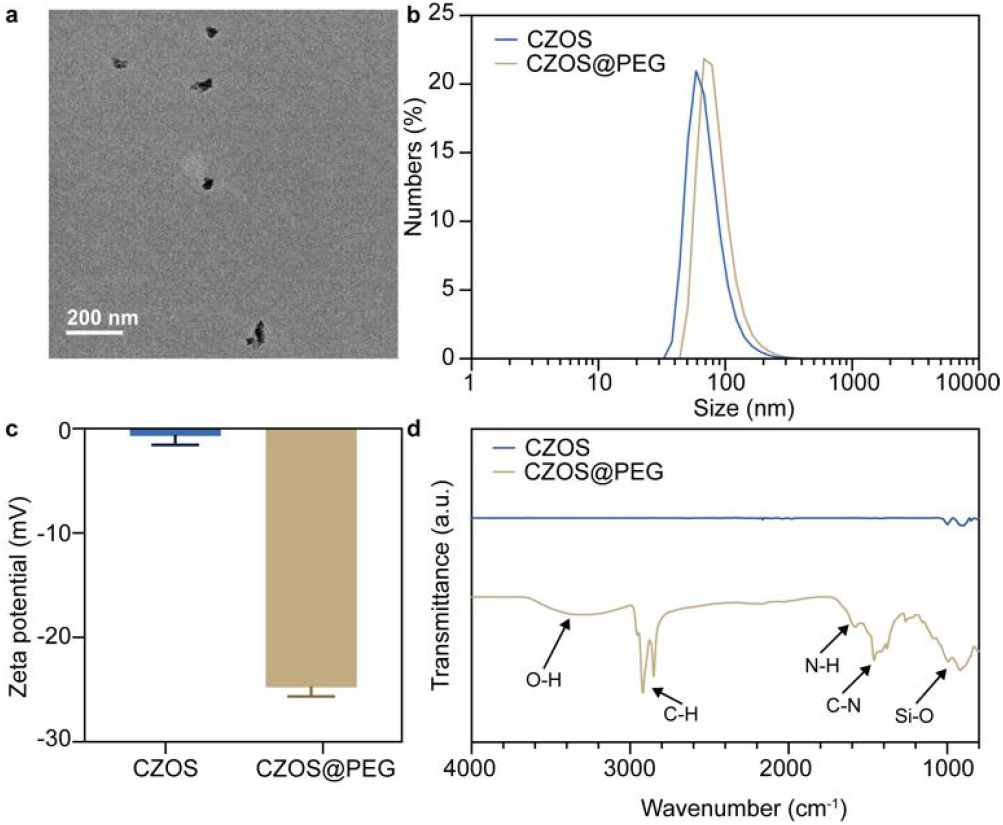
(a) Transmission electron microscopy (TEM) image of hydrophilic-modified CZOS (CZOS@PEG). (b) Hydrophilic modification-induced size variation of CZOS nanoparticles characterized by dynamic light scattering (DLS). (c) Zeta potential profiles of CZOS pre- and post-hydrophilic modification. Error bars denote standard deviations from triplicate measurements. (d) Fourier transform infrared (FTIR) spectra of CZOS pre- and post-hydrophilic modification.

Subsequently, CZOS@PEG was co-cultured with the murine macrophage cell line RAW264.7. Utilizing the endocytic effect of macrophages, CZOS@PEG-labeled macrophages were obtained, forming the biomimetic living cell probe FMN. First, the cytotoxicity of CZOS@PEG was assessed in the macrophage cell line using the Cell Counting Kit-8 (CCK-8) assay. Cells were co-incubated with a series of CZOS@PEG concentrations (0 μg mL^-1^ to 400 μg mL^-1^) at 37 °C and 5% CO_2_ for 24 h before viability assessment. No significant difference in cell growth trend was observed, indicating that FMN did not induce significant cytotoxicity (Fig. S1, Supporting Information).

The collected FMN was then immobilized within the aforementioned cylindrical phantom channel (diameter: 0.5 mm) using gelatin, and pulsed ultrasound was applied to evaluate the imaging performance (Fig. 6a and 6c). While keeping the CZOS@PEG concentration loaded in FMN constant (5 mg mL^-1^), it was observed that PL could not provide accurate spatial location information for FMN concentrations as low as 50 cells μL^−1^ (Fig. 6a). In contrast, USML generated by FUS excitation could further lower the necessary FMN concentration threshold, enabling accurate localization of FMN at concentrations as low as 10 cells μL^-1^ (Fig. 6a).

**Figure 6.**
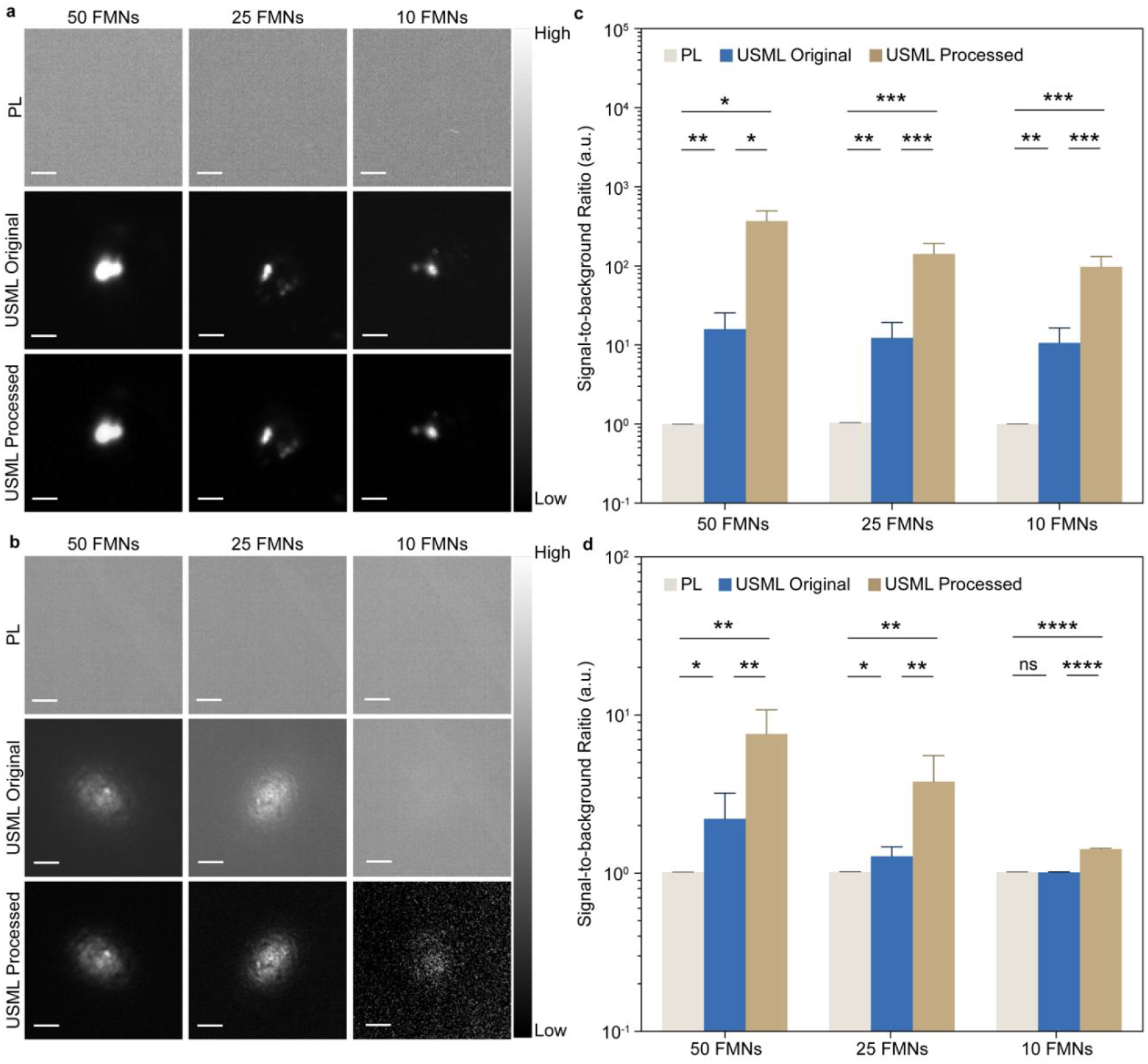
PL and USML images pre- and post-FFIR processing of trace amounts of FMNs at (a) varying concentrations and (b) in mouse subcutaneous tissue. All images have been normalized. The scale bar was defined as 0.25 mm and 1 mm, respectively. SBR comparison of PL with USML imaging pre- and post-FFIR processing on trace amounts of FMNs at (c) varying concentrations and (d) in mouse subcutaneous tissue. Error bar stands for s. d. from n = 8 replicated samples. FUS parameters applied to FMNs: repetition frequency = 1 Hz, duty ratio = 15%, and a 1500 nm long-pass filter was used to exclude excitation.

Quantitative analysis results showed that for FMN concentrations of 50, 25, and 10 FMNs μL^−1^, USML imaging increased the SBR by factors of 15.99, 11.90, and 10.67, respectively, compared to PL imaging (Fig. 6c). Furthermore, the imaging resolution of the obtained USML images could be further enhanced after FFIR processing. The SBR for imaging 50 and 25 cells μL^-1^ FMN reached 366.51 and 140.47, respectively, after FFIR. When the FMN concentration decreased further to 10 cells μL^-1^, the SBR of its USML image could still be improved from 10.46 to 96.87 after FFIR processing, significantly enhancing imaging resolution (Fig. 6c). Therefore, these results preliminarily verified the feasibility of the designed imaging system for locating trace samples.

Building on this, to explore whether the system could maintain high-sensitivity tracking imaging within deep tissues in vivo, trace amounts of FMN were used as targets for imaging observation under the coverage of mouse skin tissue (Fig. 6b and 6d). Imaging results showed that PL was ineffective for marking trace targets under tissue coverage (Fig. 6b), failing to visualize FMN at 50 cells μL^-1^ and below. In contrast, USML, benefiting from the avoidance of excitation light interference and coupled with the FFIR processing pipeline established based on its luminescent characteristics, enabled accurate localization of subcutaneous tissue signals (Fig. 6b). Quantitative analysis indicated that under mouse skin tissue coverage, the SBR of USML signal spots was already somewhat improved compared to the PL group. After further optimization by FFIR, the image quality was significantly enhanced. The SBR of USML signal spots generated by a minimum of 10 cells μL^-1^ FMN could still be increased from 1.02 to 1.43, thus demonstrating feasibility for high-sensitivity imaging (Fig. 6d).

### 2.4 High-Sensitivity Dynamic Monitoring of the Tumor Microenvironment

Following the in vitro confirmation of the system’s feasibility for detecting trace samples, its capability for monitoring dynamic changes within the tumor microenvironment during the tumor proliferation stage was evaluated in vivo. Studies have reported that macrophage infiltration through the basement membrane marks early malignant transformation of tumors. Significant interactions exist among tumor cells, macrophages, and blood vessels, where macrophages assist early tumor angiogenesis to enhance invasiveness^24-26^. This notion is further supported by observations of substantial vessel-associated macrophage infiltration in breast tumors^27^. High-resolution real-time imaging of the mammary gland revealed that ductal macrophages continuously extend and retract their dendrites on a cycle of approximately 2 h, surveying the entire ductal epithelium ^28^. To this end, an early-stage mammary tumor model was established in mice. FMN was administered, and its accumulation within tumor tissue was observed (Fig. 3.13a).

**Figure 7.**
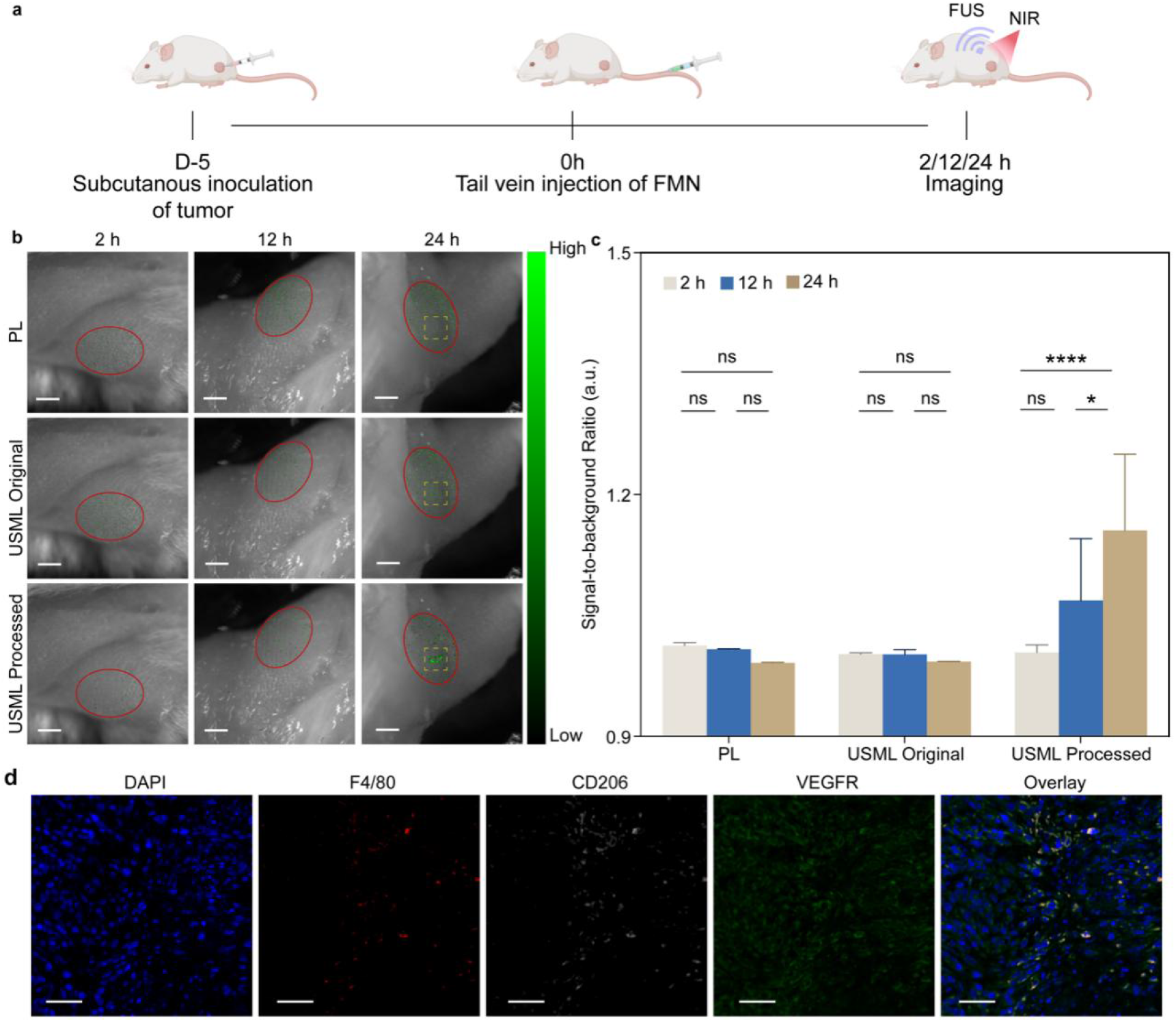
*In vivo* Imaging of FMNs. (a) Schematic illustration of the tumor microenvironment monitoring process in tumor-bearing mice. (b) PL and USML images pre- and post-FFIR processing after administration of FMNs. All images have been normalized. The scale bar was defined as 1 mm. (c) SBR comparison of PL, USML imaging pre- and post-FFIR processing at different time points. Error bar stands for s. d. from n = 4 replicated samples. FUS parameters applied to FMNs: repetition frequency = 1 Hz, duty ratio = 15%, and a 1500 nm long-pass filter was used to exclude background. (d) Immunofluorescence staining images of tumor sections from tumor-bearing mice. Scale bars were defined as 50 μm.

Monitoring signal changes within tumor tissue via PL within 24 h post-administration failed to locate accurate signals (Figs. 7b and 7c). In contrast, although USML also could not precisely localize FMN within the first 12 h, periodic luminescence was observed in the tumor tissue at 24 h. Processing the USML images with FFIR, indexed to the FUS excitation frequency (1 Hz), significantly improved imaging quality, increasing the SBR from 1.00 to 1.16 (Fig. 7c). Furthermore, FMN was observed to form relatively well-defined, concentrated clusters at the tumor site (Fig. 7b), indicating the enrichment of FMN within the early-stage tumor.

On day 7 post-FMN administration, ex vivo histological analysis of major organs and complete blood count (CBC) analysis were performed. No significant histological differences were observed between the administration group and the control group. Concurrently, no significant changes were noted in white blood cells, red blood cells, platelets, or hemoglobin levels, indicating good biocompatibility of FMN (Fig. S2 and S3). To further verify whether FMN clusters within tumor tissue, following the observation of luminescent signals post-FMN administration, tumors were rapidly excised from the tumor-bearing mice. Fluorescence histochemical staining was performed to label neovasculature and macrophage-specific markers within the tumor tissue. As shown in Fig. 7d, co-localization of fluorescence signals for the macrophage markers F4/80 and CD206 with CD309 (biomarker for vascular endothelial growth factor receptor, VEGFR) was observed. This indicates macrophage enrichment in tumor neovascular areas, which aligns with the FMN enrichment trend in tumor regions suggested by the FUS-responsive luminescence. This finding further substantiates the feasibility of using FMN for targeted labeling of early-stage tumor regions with high proliferative activity.

## 3. Conclusion

To summarize, we have constructed a macrophage-integrated USML imaging system for early-stage tumor. By harnessing the innate tumor neovascular homing ability of macrophages, active targeting of the proliferative tumor microenvironment was achieved. Subsequent excitation via FUS generated NIR-IIb USML signals, enabling high-resolution localization and dynamic tracking of trace targets. The unique acoustic-to-optical conversion property of the material circumvents the dependency on external optical excitation inherent in conventional OI modalities. This fundamental shift effectively eliminates interference from excitation light and mitigates issues related to photobleaching. Furthermore, the emission within the NIR-IIb window minimizes the impact of tissue autofluorescence, substantially improving SBR and imaging sensitivity. Consequently, precise spatial localization of signals from as few as 10 FMNs was accomplished. The post-processing technique, FFIR, effectively distinguished periodic signal components from stochastic background noise, markedly enhancing both the detection sensitivity and spatial resolution under low SBR conditions, and yielding more reliable imaging outcomes.

This work demonstrates a novel strategy that synergizes designed mechanoluminescent materials, cellular delivery vectors, and advanced computational imaging. It provides a promising platform for the high-sensitivity, high-resolution dynamic monitoring of deep-seated tumor microenvironments, addressing several limitations associated with current optical imaging techniques as outlined in the introduction.

## 4. Experimental Section

### Materials

Manganese sulfide (MnS, Mn ≥ 44%), calcium carbonate (CaCO_3_, 99.99%), and sodium citrate (99%) were purchased from Aladdin Co., Ltd. Zinc sulfide (ZnS, 99%) and Trierbium dioxide (Er_2_O_3_, 99.999%) were purchased from Adamas Co., Ltd. mPEG-Silane (98%) was purchased by Alfa Aesar Co., Ltd. SYLGARD 184 (PDMS) was purchased by Dow Corning Co., Ltd.

### Characterization

X-ray diffraction (XRD) data were collected by a Bruker D2 Phaser desktop XRD (BRUKER). Transmission electron microscopy (TEM) images were acquired with a 120 kV JEM-1400 Plus transmission electron microscope (JEOL). The emission spectra were recorded using an FX2000 spectrometer and a NIR1700 (Ideaoptics). Imaging data were performed via a near-infrared II (NIR-II) fluorescence imaging system (NIROPTICS).

### Synthesis of CaZnOS:xMn,yEr nanoparticles (CZOS)

According to the stoichiometric ratio (Ca_1-y_Zn_1-x_OS:xMn, yEr), weigh the corresponding masses of MnS, CaCO_3_, ZnS, and Er_2_O_3_ respectively. Add 10 mL of ethanol and grind the solid mixture thoroughly until a uniform suspension was formed. After the ethanol was evaporated, the above operation was repeated twice. Following the final evaporation step, the solid mixture was transferred to an alumina boat and placed into a tubular furnace. Under an argon atmosphere, the temperature was raised to 1100 °C and held for 3 hours. Upon completion of the reaction, the mixture was cooled to room temperature. The molten product was collected and further ground and crushed to obtain the powder sample.

### Synthesis of CaZnOS:xMn,yEr@mPEG-Silane (CZOS@PEG)

60 mg of the as-prepared CZOS powder was thoroughly ground and ultrasonically dispersed in 60 mL sodium citrate aqueous solution (wt 1%) to form a homogeneous dispersion. The mixed system was heated to 50 °C and maintained at this temperature for 2 h. The resulting particles were collected by centrifugation, washed twice with ethanol, and re-dispersed in an aqueous phase (1 mg·mL^-1^). Subsequently, the aqueous dispersion was co-incubated with mPEG-silane (10 mg·mL^-1^) at room temperature for 2 h with occasional shaking. After being washed twice with ethanol again, the particles were finally dispersed in ethanol for storage.

### Photoluminescent properties of CZOS

To investigate the PL properties of CZOS, 0.1 g of the powder sample was mixed with 0.3 g of PDMS in a cubic mold (1 cm×1 cm×1 cm). The mixture was heated at 75 °C for 10 h to prepare a phantom. The phantom was exposed to a CW 365 nm laser, and a 400 nm long-pass filter was used to eliminate excitation. The PL spectra of CZOS with different Mn^2+^/Er^3+^ doping ratios were measured.

### USML properties of CZOS

A cylindrical model with a diameter of 2 cm and a height of 0.5 cm was fabricated with PDMS as described in the preceding synthesis protocol. A central channel with a diameter of 0.5 mm was created in the model. The channel was filled with CZOS powder, and the surface was subsequently sealed by thermally curing an additional layer of PDMS under the same conditions to encapsulate the CZOS. The CZOS-doped phantom was fixed at the focal position above a FUS transducer, and excitation was performed at a constant duty cycle (15%) and repetition frequency (1 Hz). A 40% (w/v) gelatin aqueous solution was prepared by heating gelatin to 50 °C with rapid stirring until complete dissolution. 50 μM hemoglobin and 1 v/v% Intralipid were added to the gelatin solution, which was continuously stirred rapidly until homogeneous. After cooling to room temperature, the gelatin phantom was obtained. A series of gelatin phantoms with different thicknesses (0-5 mm) were placed over the CZOS phantom to simulate *ex vivo* tissues. The luminescence intensity of CZOS in the NIR-II window under FUS excitation with different powers was compared at various depths, using a 1500 nm long-pass filter to eliminate background interference.

### Fourier Frequency-transform Identification Reconstruction (FFIR) Algorithm

The FFIR algorithm comprises three core steps: frequency-domain transformation, feature extraction, and post-processing. The core of the algorithm lies in processing the temporal intensity variation at each pixel by transforming it into the frequency domain. For each spatial position (*x, y*) in the image sequence, its intensity variation over time, *I*_*t*_ *(x, y)*, is extracted (Eq. 4.1):

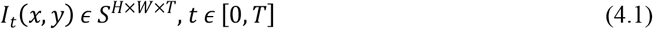

where S represents the image sequence, H and W are the height and width of a single frame, respectively, and T is the total number of frames. A Fast Fourier Transform (FFT) is applied to this temporal signal *I*_*t*_ *(x, y)*, converting it from the time domain to the frequency domain to obtain its complex-valued spectrum *F (x, y, f)* (Eq. 4.2):

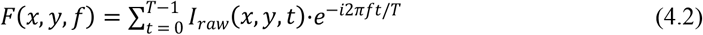

By taking the modulus of the complex result, the phase information is discarded, retaining the amplitude component to obtain the amplitude spectrum *A (x, y, f)* (Eq. 4.3):

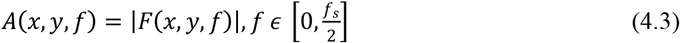

where *f*_*s*_ is the sampling frequency of the amplitude spectrum. The amplitude value *A (x, y, f)* directly represents the energy contribution of the frequency component *f* in the original signal, reflecting the signal strength at that frequency.

Frequency-domain analysis results indicate that excitation signals with periodic characteristics form distinct peaks at the target frequency (fundamental frequency) and its integer multiples (harmonics, e.g., second, third harmonics). Better periodicity of the data leads to more pronounced peaks at the fundamental frequency and harmonics (Fig. S4a). This spectral feature reveals the presence of a nonlinear response during excitation. In contrast, background noise exhibits random, broadband distribution characteristics in the time dimension, manifested as typical broadband features without harmonic structure in the frequency domain (Fig. S4b). At 1 Hz, the intensity difference between signal pixels and noise pixels were observed to be approximately 10-30 fold. Imaging based on the intensity component at the target frequency can further significantly amplify this intensity difference, enhancing signal distinguishability.

Therefore, to extract the luminescence response associated with the target FUS excitation cycle frequency (*f*_*target*_ = 1 *z*), the amplitude values within a narrow frequency band centered at *f*_*target*_ (*f*_*target*_ ± Δ*f*), are integrated. The integrated amplitude within this band is defined as the quantitative intensity *O* (*x, y*) of that pixel’s response to the target frequency (Eq. 3.4):

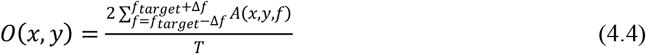

By selecting frequency components within the target frequency and tolerance band, then dividing by the total number of frames used for the FFT, the intensity value for that pixel at the specified frequency and tolerance range for a single composite image is obtained. Repeating this process for all pixels in the image generates a two-dimensional spatial distribution map *O* (*x, y*)

The initial amplitude image *O* (*x, y*) generated via frequency-domain extraction may still contain shot noise or anomalous pixels caused by random errors. To optimize image quality while preserving important spatial edge features, a two-dimensional median filter is subsequently applied for post-processing. A sliding window is used, where the intensity values of pixels within the window are sorted, and the median value is assigned as the output for the center pixel. This operation can be mathematically simplified as follows (Eq. 3.5):

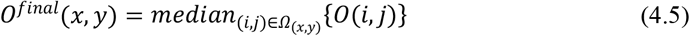

where *Ω*_(*x, y*)_ denotes the neighborhood window centered on pixel (*x, y*). This step effectively suppresses impulse noise while maximally preserving image edge sharpness, ultimately yielding a clear and reliable spatial frequency response image *O* ^*final*^ (*x, y*).

### Evaluation of Imaging Performance

In the images processed by FFIR, the signal spot area was identified. The K-means clustering was employed for segmentation, and a region of interest (ROI) with well-defined boundaries was selected to obtain the signal spot. SBR was calculated as the ratio of the average pixel intensity within the signal spot to the average pixel intensity within a background area of equal size.

### In vitro labeling and imaging of FMNs

Murine macrophage RAW 264.7 cells were purchased from Wuhan Procell Biotechnology Co., Ltd. Cells were cultured in Dulbecco’s Modified Eagle Medium (DMEM) supplemented with 10% fetal bovine serum (FBS) for 24 h. Subsequently, the cells were resuspended in complete culture medium containing 5 mg·mL^-1^ CZOS@PEG and incubated at 37 °C for 2 h. The mixture was centrifuged at 250 *g* for 3 min to collect CZOS@PEG-labeled cells (designated as FMNs). After filtering cell clusters through a 40 μm cell strainer, the FMNs were counted using a cell counter (RWD Life Science Co., Ltd.) and diluted with phosphate-buffered saline (PBS) to concentrations of 50, 25, and 10 cells·μL^-1^.

## Supporting information

12.18 Supporting Information-rev

## 5. Acknowledgment

This work was supported by the National Natural Science Foundation of China (82001945), the Natural Science Foundation of Shanghai (24ZR1452300), and the starting grant of ShanghaiTech University. The authors thank the Centre for High-resolution Electron Microscopy (ChEM), School of Physical Science and Technology, ShanghaiTech University (No. EM02161943) for the characterization support. The authors thank the Analytical Instrumentation Center (#SPST-AIC10112914), School of Physical Science and Technology, ShanghaiTech University, for the spectral test support.

## 6. Author contributions

X.Z. conceived and designed the project, supervised the overall research process, and revised the manuscript. M.Z. and J.Z. contributed equally to this work. J.Z. was responsible for the preparation and characterization of materials, as well as the design and operation of animal experiments. M.Z. mainly undertook the construction of algorithms and the processing of image data. Both J.Z. and M.Z. contributed to the writing of the manuscript. L.M., X.L., and S.X. participated in the project design and conducted part of the experimental work. All authors contributed to the general discussion and revision of the manuscript.

## 8. Data availability

All data that support the findings of this study are presented in the main text and the Supplementary Information.

