## Supplementary material for "Focused ultrasound programmed characteristic NIR-IIb lanthanide mechanoluminescence for high sensitivity bioimaging in vivo": 12.18 Supporting Information-rev

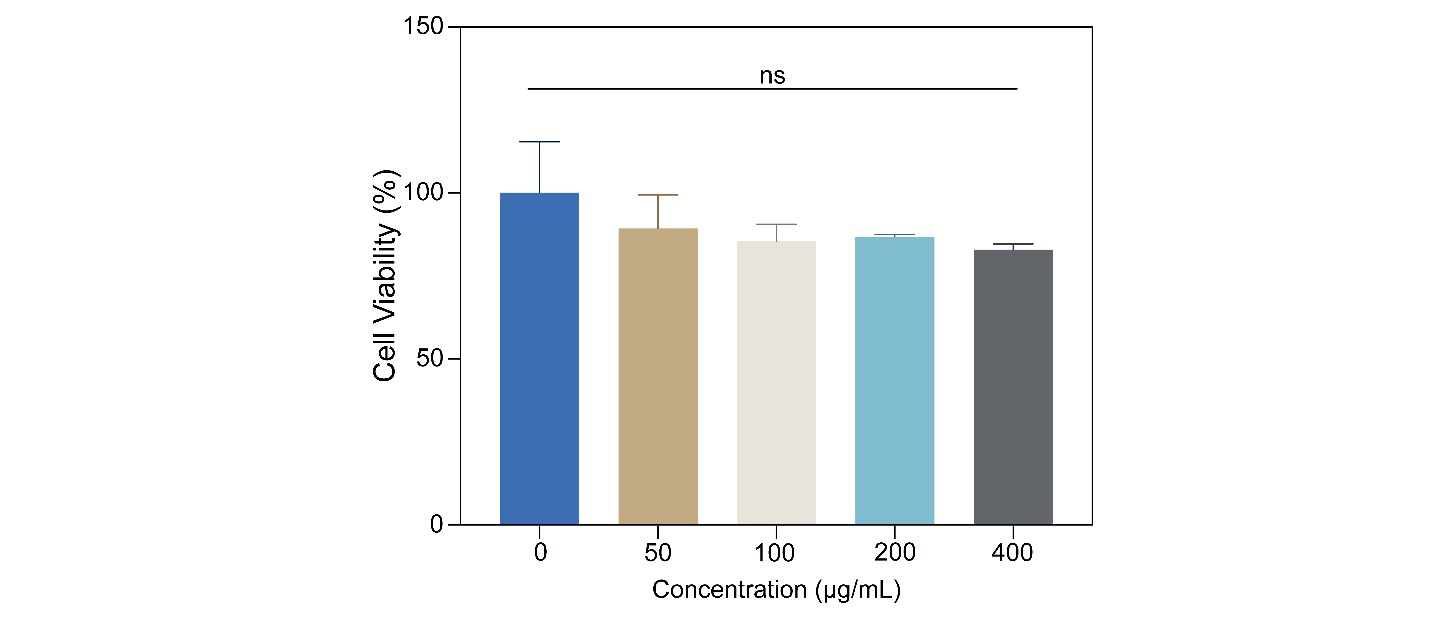

**Figure S1.** Characterizations of cytotoxicity of CZOS@PEG. Error bar stands for s. d. from n = 3 replicated samples.

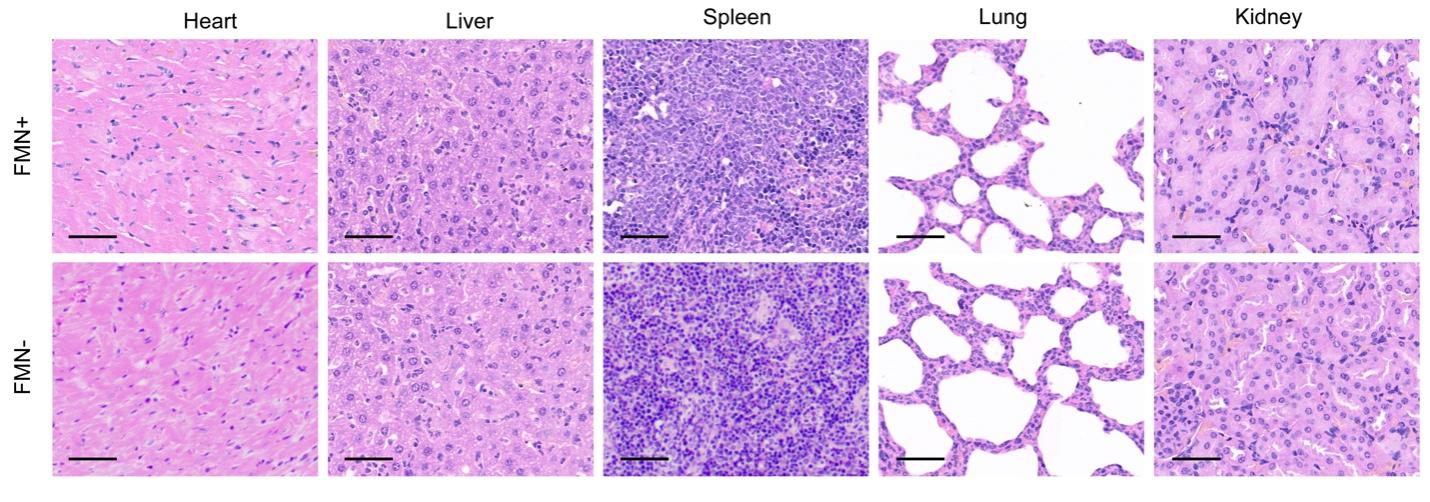

**Figure S2.** Histological analysis of heart, liver, spleen, lung, and kidney from tumor-bearing mice after FMN administration. The sections were stained with Hematoxylin and Eosin (H&E), and scale bars were defined as 50 μm.

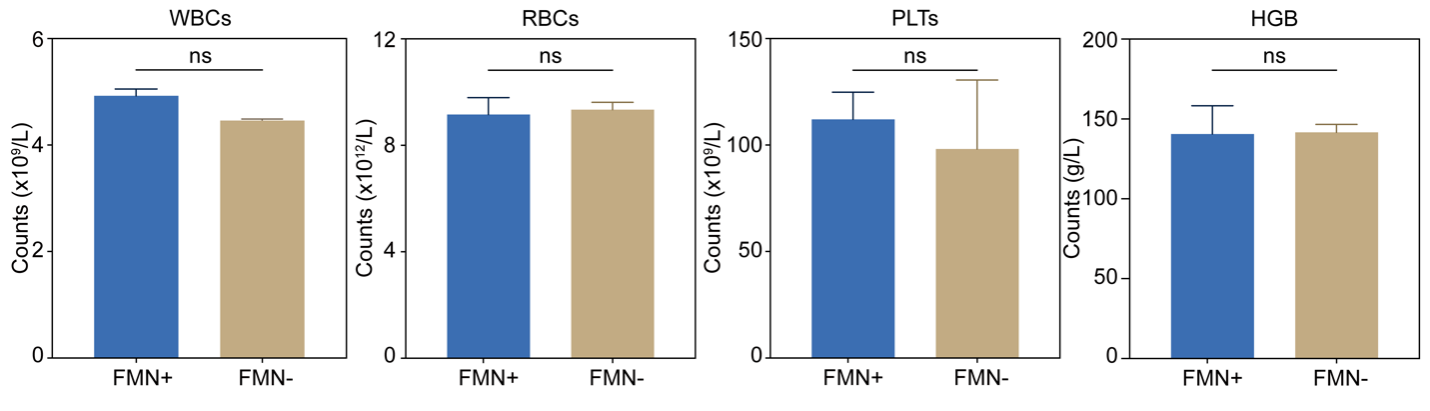

**Figure S3.** Statistical results of white blood cells (WBC), red blood cells (RBC), platelets (PLT), and hemoglobin (HGB) of tumor-bearing mice.

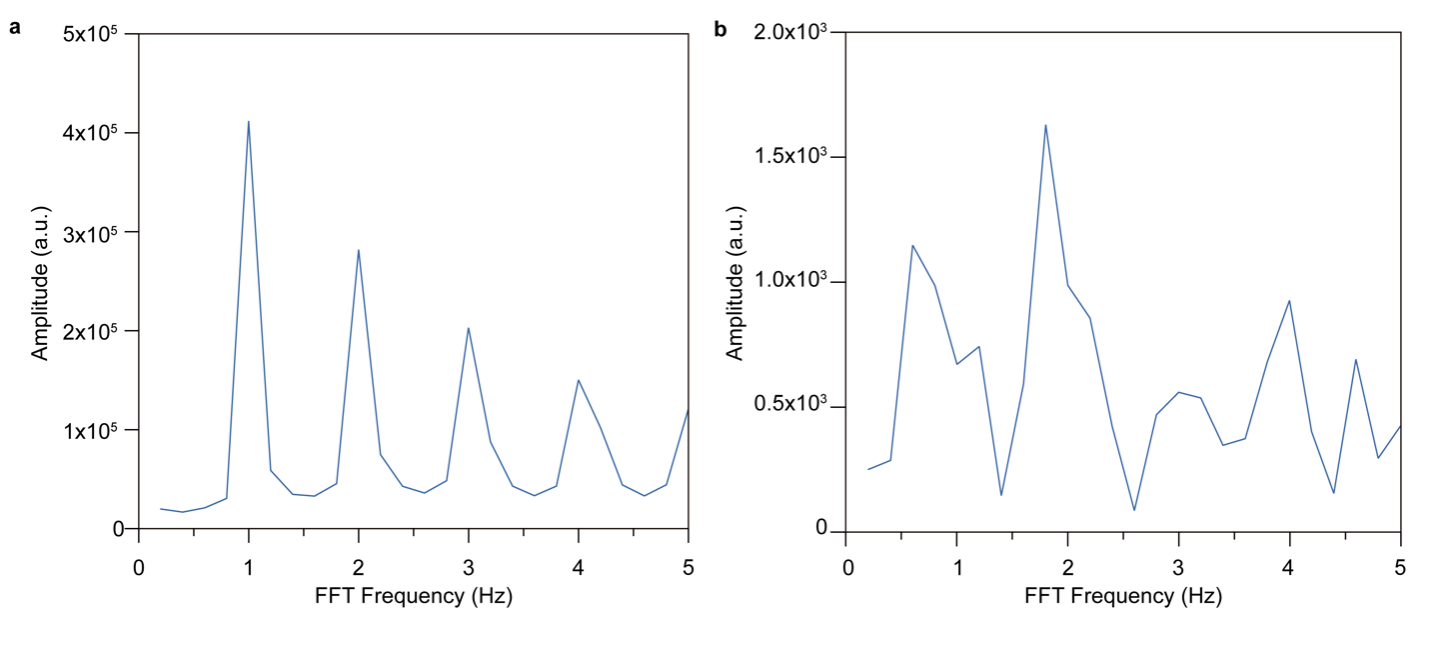

**Figure S4.** (a) Spectrum of a signal pixel after Fast Fourier Transform (FFT). (b) Spectrum of a background pixel after FFT.
